# Fanconi anemia proteins are required to maintain nucleolar homeostasis

**DOI:** 10.1101/509950

**Authors:** Anna Gueiderikh, Guillaume Rouvet, Sylvie Souquère-Besse, Sébastien Apcher, Jean-Jacques Diaz, Filippo Rosselli

## Abstract

The majority of inherited bone marrow failure (iBMF) syndromes are associated to nucleolar and/or ribosomal abnormalities, but Fanconi anemia (FA), the most common iBMF, is attributed to alterations in DNA damage responses. However, the involvement, if any, of the FA (FANC) proteins in the maintenance of nucleolar functions and/or ribosome biogenesis is yet unexplored. Here, we report that FANC pathway loss-of-function is associated to a loss of the nucleolar homeostasis, demonstrating increased rDNA rearrangements, accumulation of nucleolar DNA damage, nucleolar protein mislocalization, and a p53-independent induction of the growth inhibitory protein p21. Moreover, specifically associated to FANCA loss-of-function, which is responsible for approximately 65% of FA cases, we observed reduced rDNA transcription and rRNA processing as well as alteration in protein synthesis and polysome profiles. Thus, we have identified nucleolar consequences associated with FANC pathway deficiency, challenging current hypothesis on the physiopathology of FA.

## INTRODUCTION

The nucleolus is an extremely dynamic membrane-less nuclear substructure that is visible during the interphase, disappears at the onset of mitosis, and is reassembled at the end of telophase. Several hundreds of proteins are hosted inside the nucleolus together with the genomic sites containing the ribosomal DNA (rDNA) sequences organized in tandemly repeated clusters dispersed on the five human acrocentric chromosomes. Three major events occur in the nucleolus: the RNAPolI-mediated synthesis of the 45S/47S rRNA precursors, their processing to generate the 28S, 18S and 5.8S rRNA molecules that will be assembled with the ribosomal proteins to form the pre-ribosome units, which are successively processed to form the cytosolic 40S and 60S ribosome subunits (Boulon et al., 2010; Tsekrekou et al., 2017). In addition to its canonical role in the biogenesis of the ribosome, the nucleolus acts as a sensor of stress originating from inside the nucleolus, the nucleus or the cytosol. It can quickly lose homeostasis by modifying its activity, size, shape and protein content to adapt the cell metabolism to threithening events (Boulon et al., 2010; Tsekrekou et al., 2017). Notably, the presence of DNA damage, both inside and outside the nucleolus, constitutes a major signal leading to the nucleolar stress response characterized by rDNA transcription downregulation and mislocalization of key nucleolar proteins. The stress-induced transitory loss of nucleolar homeostasis, even in the absence of DNA damage and its signalling, is associated with the activation of the p53-p21 axis, which restrains cellular proliferation to allow time to recover from stress (Boulon et al., 2010; Tsekrekou et al., 2017).

Unrestrained, the activation of the p53-p21 axis is considered a key event responsible for the haematopoietic stem cell (HSC) attrition that characterizes inherited bone marrow failure (iBMF) syndromes, including dyskeratosis congenita (DC), Shwachman-Diamond syndrome (SDS), Diamond-Blackfan anemia (DBA) and Fanconi anemia (FA). In contrast to DC, SDS and DBA, which associate BMF and p53-p21 activation with the presence of nucleolar and/or ribosome biogenesis abnormalities (Liu and Ellis, 2006; Ruggero and Shimamura, 2014), FA is attributed to abnormalities in the DNA damage response that lead to p53-p21 activation and BMF (Ceccaldi et al., 2012; Walter et al., 2015).

FA is the most frequent and genetically heterogeneous iBMF syndrome (Bogliolo and Surralles, 2015; Ceccaldi et al., 2016; Gueiderikh et al., 2017), and is associated with developmental abnormalities, predisposition to cancer and chromosomal instability. The key function of the molecular pathway defined by FA proteins (FANC) is to resolve DNA interstrand crosslinks (ICLs) and stalled replication forks that can arise from endogeneous aldheyde metabolism or following exposure to ICL-inducing agents, as mitomycin C, cis-Pt or photoactivated psoralens (Bogliolo and Surralles, 2015; Ceccaldi et al., 2016; Gueiderikh et al., 2017; Langevin et al., 2011; Rosado et al., 2011). The FANC proteins are organized in three distinct functional groups. The FANCcore complex, which comprises FANCA, FANCB, FANCC, FANCE, FANCF, FANCG, FANCL, FAAP24, and FAAP100, is assembled to monoubiquitylate the FANCD2-FANCI (ID2) heterodimer, allowing its accumulation in subnuclear foci required for the optimal work of the third group of FANC proteins, which includes nucleases and homologous recombination proteins (Bogliolo and Surralles, 2015; Ceccaldi et al., 2016; Gueiderikh et al., 2017). Even if the loss-of-function of one among more than 20 genes is recognized as involved in FA, *FANCA* bi-allelic inactivating mutations account for more than 60% of FA cases worldwide (Gueiderikh et al., 2017). Beyond the DNA repair/DNA damage response, the FANC pathway has been associated with several other cellular and systemic biological activities, including pro-inflammatory cytokine production and responses, transcription, mRNA splicing and reactive oxygen species metabolism. However, the presence, origin and consequences of nucleolar abnormalities induced as a consequence either of a still unknown role of the FANC pathway in ribosome biogenesis or of the chronic stress state to which the FANC pathway-deficient cells are subjected has not been explored in FA.

Here, we investigated nucleolar protein localization, rDNA stability and transcription as well as translational activity in FA-deficient cells, demonstrating an involvement of the FANC pathway in the nucleolar homeostasis. Moreover, we also identified a function of FANCA in ribosome biogenesis, a possible reason accounting for the high frequency of patients with inactivating mutations in *FANCA.* We thus hypothesize that FA originates from alterations in both DNA repair and ribosome biogenesis.

## RESULTS and DISCUSSION

### The loss-of-function of the FANC pathway is associated to nucleolar proteins mis-localization

To determine the impact, if any, of the inactivation of the FANC pathway on the nucleolar homeostasis, we monitored the expression and subcellular distribution of the key nucleolar proteins fibrillarin (FBL), nucleolin (NCL) and upstream-binding nucleolar transcription factor 1 (UBF) in HeLa and U2OS cells 48h to 72h after the siRNA-mediated depletion of FANCA, FANCC or FANCD2. The depletion of any of the three FANC proteins did not impact the expression of the analysed nucleolar proteins (Fig. S1A and data not shown). In the large majority (>70%) of FANCA-proficient cells, nucleoli had irregular borders delimited by a thick ring of NCL inside which FBL and UBF are distributed in a fairly homogeneous way (Fig. 1A to D and Fig. S1B to D). In contrast, while maintaining the same quantity of nucleoli per cell (Fig. S1E), only a minority of FANCA-depleted cells (around 30%) have nucleoli with the previously described distribution of NCL, FBL and UBF and/or irregular borders (Fig. 1A to D and Fig. S1B to D). Indeed, in the majority of the FANCA-depleted cells nucleoli took either a more round and regular shape delimited by an extremely faint ring of NCL, or they appeared “empty” with the proteins displaced out of the nucleolus or eventually accumulating in nucleolar peripheral structures called “caps” (Fig. 1A to D and Fig. S1B to D). Electronic microscopy (EM) validated the occurrence in FANCA-depleted cells of less organized and/or more condensed and rounded nucleoli (Fig. 1E). The depletion of FANCC, belonging as FANCA to the FANCcore complex, increased the frequency of cells presenting nucleolar abnormalities to around 40%, a level in between that observed for WT (around 20%) and FANCA-depleted (more than 70%) HeLa cells (Fig. 1F). Even if the depletion of FANCD2, a FANCcore target, led to a general increase in the percentage of cells with an altered localization of nucleolar proteins (up to 30%), the frequency of cells with altered nucleoli reached the threshold of significance only in the cells in S/G2 (Fig. 1F).

**Figure 1:**
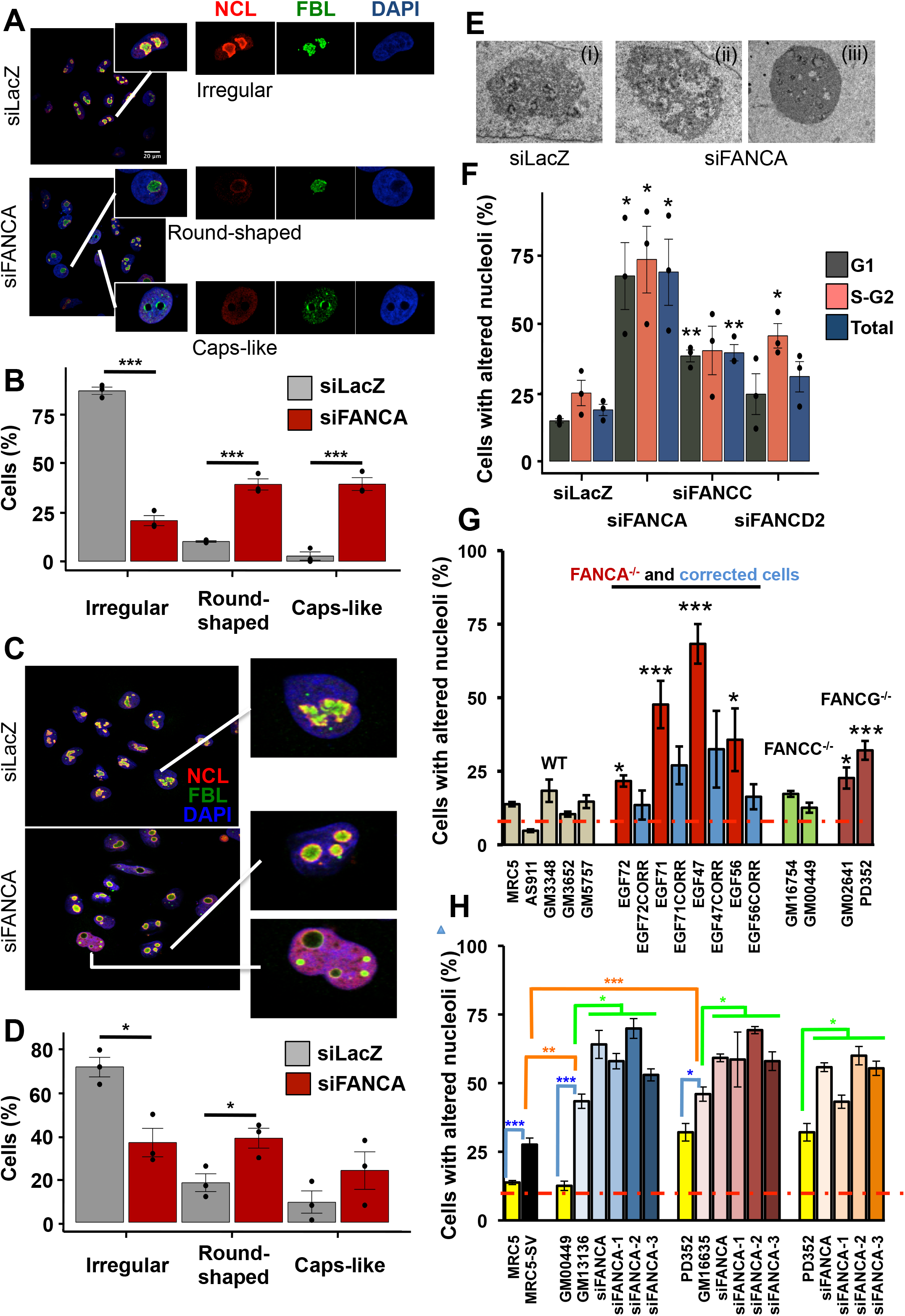
Nucleolar abnormalities in FANC pathway-deficient cells. **A and C.** Confocal microscopy images showing wide fields and single enlarged nuclei of HeLa (A) or U2OS (C) cells after transfection with untargeted (siLacZ) or FANCA-targeted siRNAs stained with antibodies against the nucleolar proteins NCL (Red) and FBL (Green), and counterstained with DAPI to visualize DNA. **B and D.** Percentage of HeLa (B) or U2OS (D) cells presenting the canonical nucleolar morphology (irregular shape) or a round or cap-like shape. Cells were transfected and stained as in A and C. Bars represent the mean of 3 independent experiments +/-sem. **E.** Electronic micrographs showing nucleolar morphology as observed in the majority of FANCA-proficient HeLa cells (i) and the disorganized or more condensed nucleoli observed 72h following siRNA-mediated FANCA depletion (ii and iii). **F.** Percentage of HeLa cells with altered nucleoli, *i.e*., round + caps-like phenotypes, 72 h following transfection with untargeted (siLacZ) siRNA or FANCA-, FANCC-or FANCD2-targeted siRNAs as a function of their cell cycle position at the moment of analysis. Cells were stained as in **A** and with anti-Cyclin A2 to discriminate G1 from S/G2 cells. **G.** Percentage of cells with altered nucleolar morphology in FANC pathway-proficient, FANCA^−/-^ and FANCA-corrected, FANCC^−/-^ and FANCG^−/-^ primary fibroblasts. Bars represent the mean of 3 independent experiments +/-sem. The dotted red line represents the mean of the WT cell lines analysed. For statistical analysis, the mean of each FANCA-deficient cell line was compared to the mean of the 5 control cells. **H.** Percentage of cells with altered nucleolar morphology in FANC pathway-proficient MRC5 primary (yellow bar) and MRC5-SV immortalized (black bar) cells, in FANCC^−/-^ primary fibroblasts (GM00449, yellow bar) and their immortalized counterpart (GM13136, light blue), FANCG^−/-^ primary fibroblasts (PD352, yellow bar) and their immortalized counterpart (GM16635, pink bar) and in FANCC^−/-^ (GM13136) and FANCG^−/-^ (GM16635) immortalized cells as well as in FANCG^−/-^ primary (PD352) fibroblasts following siRNA-mediated FANCA depletion. Bars represent the mean of 3 independent experiments +/-sem. As in G, the dotted red line represents the mean of the primary WT cell lines analysed. For statistical purpose we compared primary vs immortalized cells (blue lines), FANC-proficient vs FANC-deficient immortalized cells (orange lines) or FANC-deficient vs FANC-deficient/FANCA-depleted cells (green lines). In all panels, statistical significance was assessed with two-tailed unpaired Student’s *t*-tests by using GraphPad on-line software (*p<0.05, **p<0.01, ***p<0.005).

To further extend and validate our observations, we analysed several human primary fibroblastic cell lines from healthy and FA donors. In the five FANC pathway-proficient cell lines analysed 12% of the cells presented a non-canonical nucleolar distribution of NCL, FBL and/or UBF, with frequency spanning between 5% and 18% (Fig. 1G). With respect to mean value, all analysed FANCA-mutated cell lines presented a significant increase in the frequency of cells with altered nucleoli, frequency that is invariably reduced by the ectopic expression of the WT FANCA gene (Fig. 1G). The previous observations in primary cells isolated from FA patients support that the data obtained in HeLa or U2OS cells were not due to off-target effects of the siRNAs transfection (Fig. S1C and D). Compared to the WT cells, the two analyzed FANCC^−/-^ cell lines were indistinguishable whereas in both FANCG-deficient fibroblasts the frequency of cells with nucleolar abnormalities was significantly more elevated (Fig. 1G). Fibroblasts immortalization increases the frequency of cells with nucleolar abnormalities independently of their FANC pathway status (Fig. 1H). However, compared to the FANC pathway-proficient SV40 transformed SV-MRC5 cells, both immortalized FANCC^−/-^ and FANCG^−/-^ cell lines demonstrated a significant increase in the frequency of cells with mislocalized nucleolar proteins (Fig. 1H). Moreover, FANCA-depletion in FANCC^−/-^ or FANCG^−/-^ primary or immortalized fibroblasts led to an increase in cells with nucleolar abnormalities (Fig. 1H). Finally, nucleolar abnormalities were also observed in primary fibroblasts from a Fanca^−/-^ mouse model (Fig. S1F to H).

Overall, our observations, obtained from analysis on several cell models issued from humans and mice, unveil that the loss-of-function of the proximal part of the FANC pathway is associated to nucleolar abnormalities. Moreover, we noticed a stronger effect on the localization of nucleolar proteins when FANCA was depleted. Thus, FANCA could have both FANC pathway-associated and FANC pathway-independent roles in maintaining the nucleolar homeostasis. Homeostasis that can be challenged by stresses originating from either inside or outside the nucleus, including DNA damage, perturbations in replication and transcription, as well as abnormalities in ribosome biogenesis and in translation.

### Nucleolar stress in FA cells is associated to DNA damage but independent of ATM/ATR signaling

The existence of nucleolar anomalies associated with the loss of function of FANCA, FANCC, FANCG or FANCD2 raises the question of whether the FANC pathway maintains the nucleolar homeostasis by contrasting an endogenous stress linked to normal cellular metabolism or if its inactivation causes a stress that is the real, direct, responsible for the observed nucleolar anomalies. In light of the canonical involvement of the FANC pathway in DNA repair and replication rescue (Bogliolo and Surralles, 2015; Ceccaldi et al., 2016; Gueiderikh et al., 2017), in coordinate replication and transcription (Schwab et al., 2015) as well as in fragile site maintenance (Guervilly et al., 2015; Howlett et al., 2005; Naim and Rosselli, 2009a, b; Naim et al., 2013), we wondered if the nucleolar abnormalities observed in FANC pathway-deficient cells were an outcome of increased rDNA damage and/or instability or a result of increased DNA damage signaling.

rDNAs are considered chromosomal fragile sites and recombination hot-spots frequently rearranged in cancer as well as in DNA repair-deficient syndromes (Killen et al., 2009; Larsen and Stucki, 2016; Stults et al., 2009; Warmerdam et al., 2016). In accord with the role of the FANC pathway in genome stability maintenance by avoiding DNA damage or its accumulation, depletion of FANCA or FANCD2 led to both an increased level of cells with nucleoli presenting γ-H2AX foci, readout of the presence of breaks in rDNA, and the induction of rDNA rearrangements (Fig. 2A and B). Notably, transcription inhibition by exposure to ActD, a recognized inducer of nucleolar stress, failed to increase the frequency of cells with γ-H2AX-stained nucleoli (Fig. 2A), suggesting that the γ-H2AX foci observed in FANCA-or FANCD2-depleted cells likely reflect unrepaired or mis-repaired DNA breaks due to alterations in stalled replication fork rescue. According to Schwab and collaborators (Schwab et al., 2015), the loss of coordination between replication and transcription, that in FA is associated to an accumulation of RNA-DNA hybrid structures or R-loops (Garcia-Rubio et al., 2015), is the main cause of the spontaneous genome instability observed in FANC pathway deficient cells. Thus, R-loops accumulation in FANC pathway-deficient cells reflects replication forks delay or stalling, events that lead to their collapse and to the induction of one-handed DNA double-strand breaks. Similarly to genome-wide data previously published (Garcia-Rubio et al., 2015), we observed that the level of R-loops inside the nucleolus is significantly increased by FANCA or FANCD2 depletion (Fig. 2C and 2D and Fig. S2A and B). In accord with a direct requirement of FANCD2 in the removal of R-loops from rRNA coding sequences downstream the FANCcore complex, ChIP-qPCR analysis demonstrated that the association of FANCD2 with these regions is lowered following FANCA depletion (Fig. 2E). Unfortunately, even if transcription inhibition restored replication and genome breakage in FA cells (Schwab et al), it cannot be used to recover their nucleolar abnormalities, because it is *per se* a major source of nucleolar stress.

**Figure 2:**
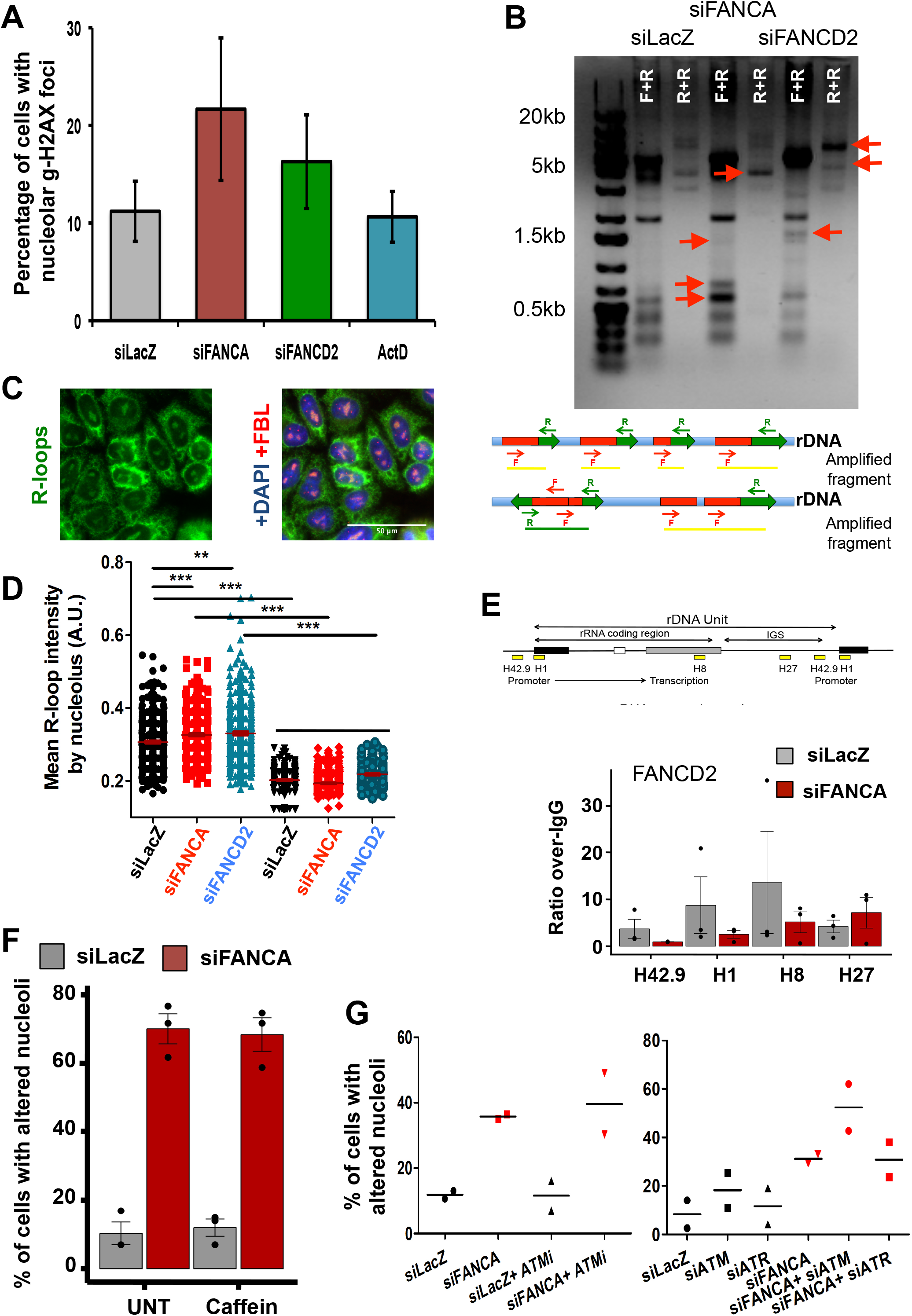
Nucleolar fragility and DNA damage signalling in FANCA-deficient cells. **A.** Percentage of HCT116 cells with γH2AX foci-positive nucleoli. The cells were transfected with the indicated siRNAs and analysed 72 h later or treated with the transcription inhibitor actinomycin D (ActD) (10 ng/mL/16 h). **B.** Genomic PCR analysis of rDNA units 72h following FANCA or FANCD2 depletion in HeLa cells by using forward(F)-reverse(R) or reverse-reverse primers (see Inset). Red arrows indicate new bands appearing in FANC-depleted cells as a consequence of genomic rearrangements in rDNA regions. **C and D.** Immunofluorescence microscopy of HeLa cells co-stained with anti-R-loops (green), anti-FBL (red) and DAPI (blue) to evaluate the nucleolar levels of DNA-RNA hybrids. Dots in the diagram represent the intensity of R-loop-staining measured using CellProfiler software with FBL staining as a nucleolar marker for each cell 48 h after transfection with untargeted, FANCA-targeted or FANCD2-targeted siRNA. The cells were incubated with RNaseH, which specifically eliminates DNA-RNA hybrids, to validate the antibody specificity. The red horizontal lines represent the mean value. A representative experiment of 3 is shown. At least 100 cells were scored for each condition. Statistical significance was assessed with a Z (normal distribution) test (***p<0.005). **E.** Top, simplified diagram showing the organization of one rDNA repeat. Yellow boxes indicate the regions amplified by ChIP-qPCR analysis: H42.9 is localized in the promoter, H1 at the start codon, H8 in the final region of the RNAPolI-transcribed region, and H27 in the inter-rDNA gene sequence (IGS), which is not transcribed by RNAPolI. Bottom, distribution of FANCD2 on the rDNAs of HeLa cells, as determined by ChIP-qPCR analysis. ChIP was performed 48h after transfection with the indicated siRNAs. Bars represent the mean of three experiments +/-sem. **F.** Percentage of HeLa cells with altered nucleoli following transfection with untargeted or FANCA-targeted siRNAs. Cells were exposed to diluent or caffeine, a widely used ATM/ATR-signalling inhibitor, 36 h before fixation. Bars represent the mean of three independent experiments. **G.** Left, diagram showing the percentage of cells with altered nucleoli in HeLa cells transfected with untargeted or FANCA-targeted siRNAs and treated or not with an ATM-specific inhibitor (ATMi). Right, diagram showing the percentage of cells with altered nucleoli in HeLa cells transfected with untargeted, FANCA-, ATM-, ATR-, FANCA+ATM-or FANCA+ATR-targeted siRNAs. Dots represent the value obtained in each realized experiment; lines indicate the mean of two experiments.

DNA breakage-induced H2AX phosphorylation is mediated by ATM signalling that is (a) constitutively active in FANC pathway-deficient cells (Guervilly et al., 2008; Kennedy et al., 2007), and (b) involved, at least in response to ionizing radiation-induced DNA damage, in the displacement of the nucleolar proteins in the nucleoplasm where some of them, including NCL and nucleophosmin (NPM1 or B23, which is also displaced in FANCA-deficient cells (Fig. 3B)), participate to the DNA damage response (Kidiyoor et al., 2016; Kruhlak et al., 2007; Ma and Pederson, 2013; van Sluis and McStay, 2015; Warmerdam et al., 2016). Thus, we asked if the nucleolar proteins displacement observed in FANC pathway-deficient cells was dependent on the activation of ATM or its related kinase, ATR. Inhibition or depletion of ATM and/or ATR, failed to rescue the nucleolar proteins mislocalization in FANCA-deficient cells, demonstrating that nucleolar abnormalities were not consequence of unrestrained ATM/ATR signalling (Fig. 2F and G).

**Figure 3:**
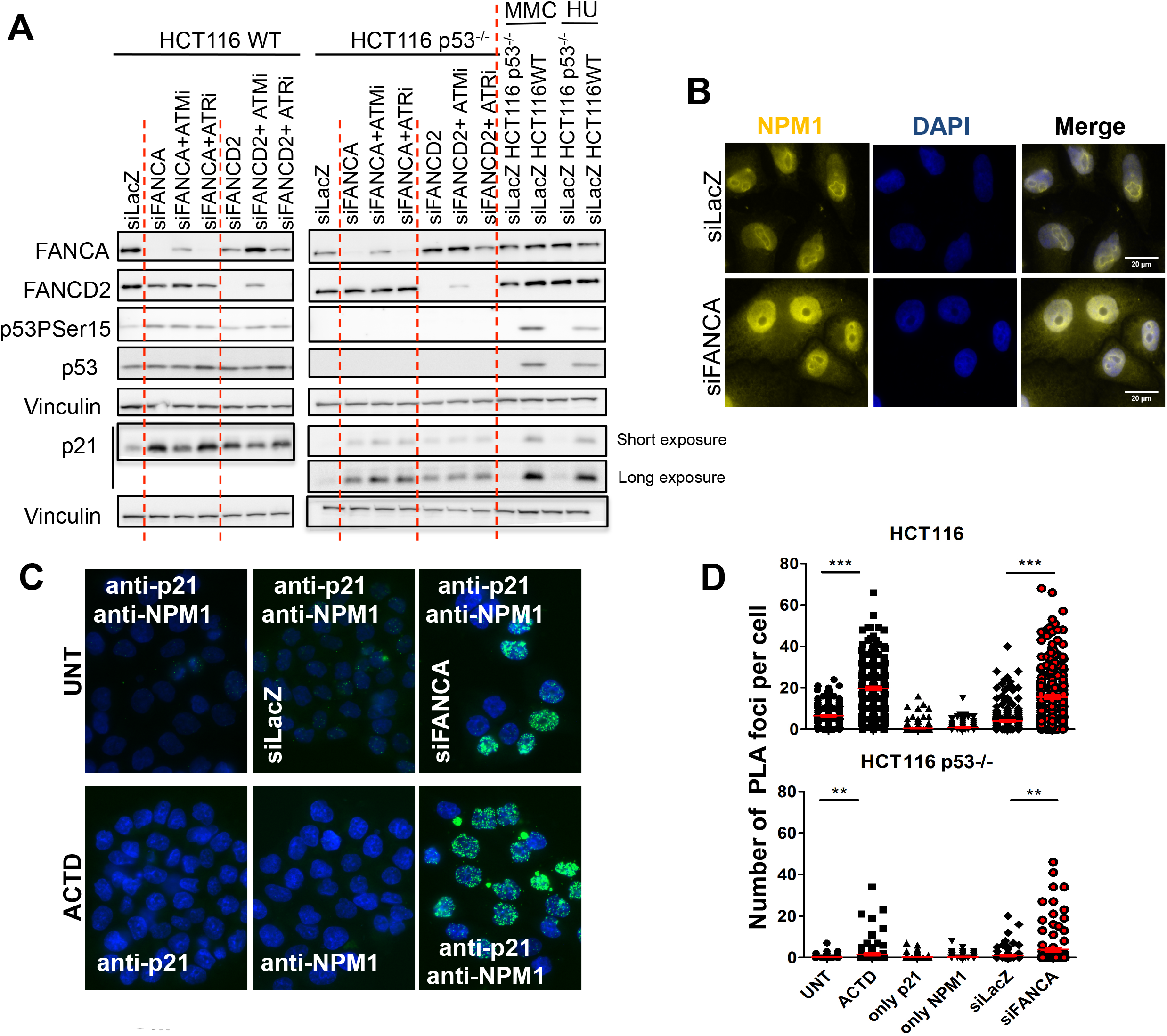
FANC pathway inactivation leads to p53-independent p21 induction. **A.** Representative Western blots showing the expression of the indicated proteins in HCT116 or HCT116-p53^−/-^ cells transfected with untargeted (siLacZ) or targeted (FANCA or FANCD2) siRNAs and treated or not with ATM or ATR inhibitors. The response to two known genotoxic agents, mitomycin C (MMC) and hydroxyurea (HU), is also shown. **B.** Representative images of wide fields of HeLa cells showing the cellular distribution of NPM1 (yellow) as a function of FANCA expression. Nuclei were stained with DAPI. **C.** Representative images of PLA on untransfected HCT116 cells or HCT116 cells 72 h following transfection with the indicated siRNAs. Untreated and ActD-treated samples were stained with anti-p21 and/or anti-NPM1 antibodies. **D.** Diagrams showing the quantitave analysis of a representative experiment. PLA foci were quantified on epifluorescence images using CellProfiler software in individual HCT116 or HCT116-p53^−/-^ cells. Statistical significance was assessed with the Z (normal distribution) test (**p<0.01, ***p<0.005).

Overall, the above data support the hypothesis that in FANC pathway deficient cells the loss of coordination between replication and transcription (Schwab et al., 2015), having an impact on the elimination of the R-loops and the progression of the replication forks, leads to DNA breaks, which, in turn, as reported in response to any other exogenous sources of DNA damage, are responsible for both the rDNA rearrangements and the re-localization of the nucleolar proteins outside the nucleolus. Since the absence of a functional FANC pathway directly leads to breaks into the rDNA, ATM/ATR signalling is not required to reorganize nucleolar structure and activity as instead observed when DNA damage is induced outside the rDNAs.

### FANC pathway deficiency leads to p53-independent p21 stabilization

Given above, it appears that the FANC pathway participates in the maintenance of the nucleolar homeostasis mainly by alleviating spontaneous stress resulting from replication/transcription conflicts. Thus, the question arises whether the observed nucleolar stress is simply a new feature of FA cells or if, on the contrary, it plays a role in the FA phenotype. The induction of the growth inhibitor protein p21 is one of the key events induced both in response to DNA damage downstream the ATM/ATR-mediated phosphorylation of p53 than during the nucleolar stress response, mediated by both p53-dependent and -independent mechanisms. Interestingly, the aberrantly constitutive induction of the p53-p21 axis has been considered as a key pathological event in FA, mainly responsible for the HSC attrition that leads to BMF of the syndrome (Ceccaldi et al., 2012; Walter et al., 2015). Consequently, we wondered if the loss of nucleolar homeostasis we observed in FANC-deficient cells was involved in p21 induction. Expectedly, FANCA or FANCD2 depletion increased p53 and/or P-p53 and p21 levels in HCT116-p53-proficient cells (Fig. 3A, left). Interestingly, in FANCA-or FANCD2-deficient cells, ATM or ATR inhibition did not have major effect on p53, even if p21 expression appeared lowered following ATM inhibition (Fig. 3A, left). Then, we looked at p21 expression in a p53-KO genetic background. As reported (Fig. 3A), FANCA or FANCD2 depletion was still associated to an increased level of p21 also in absence of p53, suggesting that p21 expression in FA depends on both DNA damage, which induces p21 rigorously in p53-dependent manner (Fig. 3A, right), and nucleolar stress-dependent activities.

It has been proposed that p21 level can be stabilized in a p53-independent manner by an interaction with the nucleolar protein NMP1 (Xiao et al., 2009) that, as observed for NCL or FBL, is also largely translocated from the nucleolus in the nucleoplasm in FANC proteins depleted cells (Fig. 3B). Given the previous observations, we wondered if the observed increased level of p21 was due to it interaction with NPM1. Thus, we looked at the co-localization of NPM1 and p21 in FANC pathway-depleted cells by using the proximity ligation assay (PLA). We demonstrated a strong nuclear colocalization of NPM1 and p21 in FANCA-or FANCD2-depleted cells (Fig. 3C and D and data not shown). However, we cannot provide the formal proof that p21 stabilization relies to NPM1 interaction. Indeed, unfortunately, NPM1 depletion *per se* activates p53-dependent and -independent pathways leading to p21 induction/stabilization, masking its consequences when FANCA was co-depleted (Fig. S2C). Moreover, interfering with the interpretation of the results, co-depletion of FANCA or FANCD2 with NPM1 was found to be extremely toxic in the cells we used.

However, even if we cannot formally demonstrate that p21 stabilization in FA cells relies to its interaction with NPM1, it is clear that p21 increase in FA takes place in both a p53-and ATM/ATR-dependent manner, as a consequence of the accumulation of DNA damage, and in a p53-and ATM/ATR-independent manner, as likely consequence of the observed nucleolar stress.

### FANCA deficiency is associated with reduced rDNA transcription

To further extend our observations and shed light on the relationship between the FANC pathway ad the nucleolar homeostasis, we wanted to determine its involvement, if any, in the process of rRNAs production, one of the main nucleolar activities. Pulse-chase experiments with ^32^P-orthophosphate followed by electrophoretic separation of the pre-rRNA species revealed that the 45S/47S rRNA precursors are produced at a lower rate in FANCA-deficient than in FANCA-proficient cells and that their processing, although qualitatively unaltered, is slowed down in absence of FANCA (compare 28S and 18S levels in siLacZ vs siFANCA) (Fig. 4A). Importantly, FANCD2 deficiency did not led to similar alterations in rDNA transcription or rRNA processing (Fig. S2D), supporting the possibility of a different requirement for FANCA in nucleolar homeostasis.

**Figure 4:**
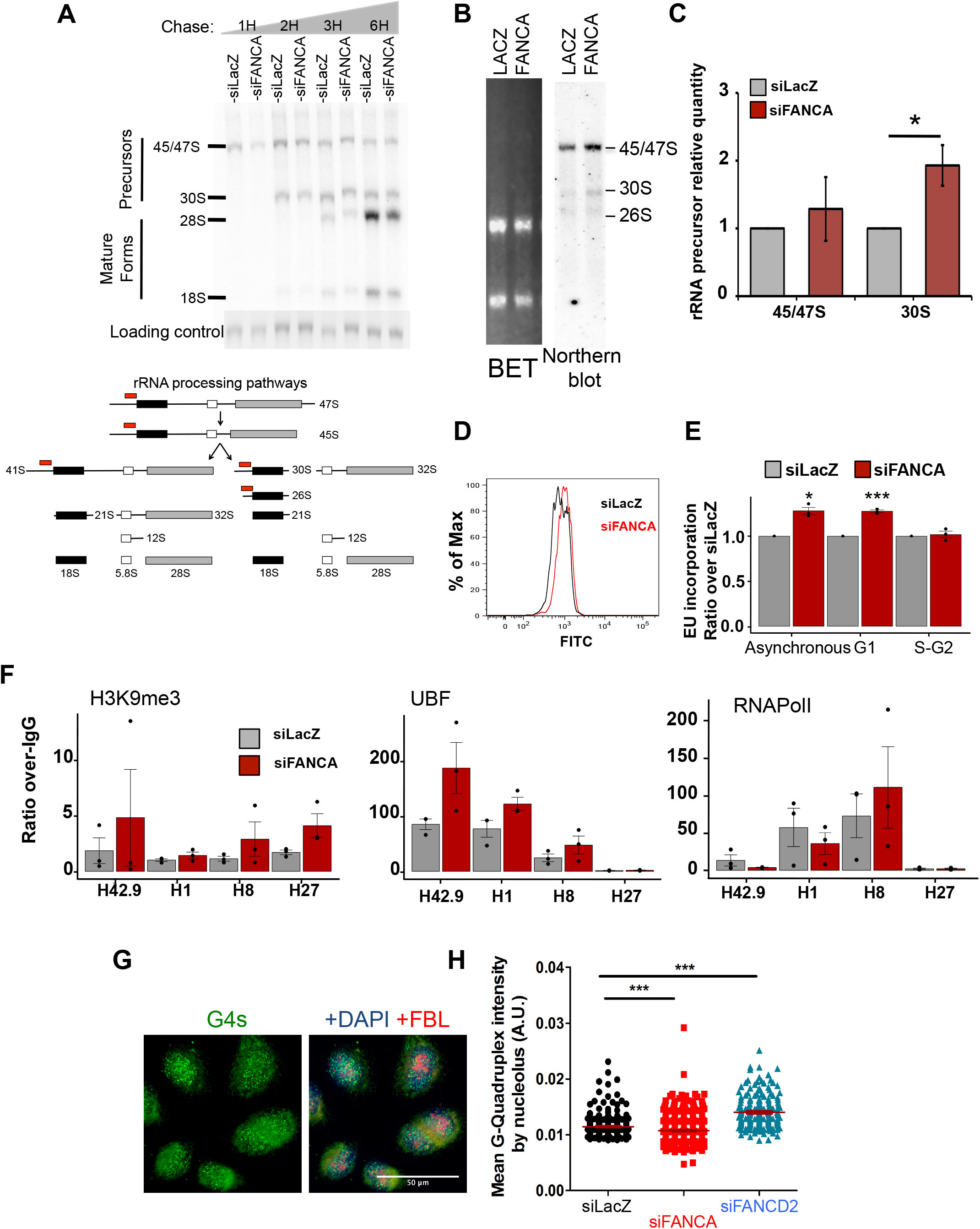
rDNA transcription and rRNA processing abnormalities in FANCA-deficient cells. **A.** Top, representative experiment showing precursor and mature rRNAs. After 72 h of siRNA transfection, HeLa cells were labelled for 20 min with ^32^P-orthophosphate and chased with cold orthophosphate for 1, 2, 3 and 6 h. The EtBr-stained gel is shown at the bottom as a loading control. Bottom, simplified diagram showing two major pre-rRNA processing paths. Names of the intermediates are given according to the literature. Red boxes indicate the probes used in the Northern blot analysis presented in C. **B.** Northern blot analysis performed with the probe indicated in A. RNAs were isolated 72 h after siRNA transfection of HeLa cells. The EtBr-stained gel is shown as a loading control. A representative experiment is shown. **C.** Quantification of four independent experiments performed as in B. The quantity measured in FANCA-depleted cells was adjusted to that in FANCA-proficient cells, which was set to 1. Bars represent the mean of 4 independent experiments +/-sem. Statistical significance was assessed with one-tailed Student’s *t*-tests (*p<0.05). **D.** Example of FACS analysis measuring the level of EU incorporation into cells following 15 min of incubation. The black line represents the profile of HeLa cells transfected with an untargeted siRNA, and the red line shows the profile of HeLa cells 72 h following transfection with siRNAs targeting FANCA. **E.** Quantification of the EU incorporation in asynchronous cells (total) and as a function of the cell cycle phase of the cells assessed with cyclin staining (G1 vs S-G2) 72 h after transfection. The absolute value (mean) in FANCA-depleted cells was compared to that in FANCA-proficient cells, which was set to 1. Bars represent the mean of 3 independent experiments +/-sem. Statistical significance was assessed with two-tailed Student’s *t*-tests (***p<0.005). **F.** Distribution of H3K9me3, UBF and RNAPolI on the rDNAs of HeLa cells, as determined by ChIP-Q-PCR analysis. ChIP was performed 48 h after transfection with the indicated siRNAs. Bars represent the mean of three experiments +/-sem. **G and H.** Immunofluorescence microscopy of cells co-stained with anti-G4 (green), anti-FBL (red) and DAPI (blue) to evaluate the nucleolar levels of G4 structures. Dots in the diagram represent the intensity of G4-staining measured in epifluorescence images using CellProfiler software with FBL staining as the nucleolar marker for each cell 48 h after transfection with untargeted, FANCA-targeted or FANCD2-targeted siRNAs. Horizontal lines represent the mean value. At least 100 cells were scored for each condition. Statistical significance was assessed with a Z (normal distribution) test (***p<0.005).

Northern blot analysis allowed us to validate that FANCA depletion leads to increased levels of both 45S/47S and 30S rRNA precursors (Fig. 4B and C), supporting a delay in pre-rRNA processing, in line with our previously published observations indicating lower levels of 28S in FA cells (Zanier et al., 2004). By FACS analysis, we quantified nascent RNA synthesis by measuring the nucleolar incorporation of the modified RNA precursor 5-ethynyl uridine (EU) (Jao and Salic, 2008), which demonstrated that FANCA depletion resulted in a slight but consistent increase in the incorporation of EU, mainly in G1 cells (Fig. 4D and E). Next, by ChIP-qPCR analysis, we observed that FANCA depletion resulted in an enrichment on the rDNA promoter region of both UBF and the H3K9me3 histone mark, which is associated with actively transcribed chromatin, whereas RNAPolI accumulated on the distal parts of the rDNA (Fig. 4F). Thus, the ChIP-qPCR data are consistent with an enhanced rate of transcription initiation and a delay in transcription completion, respectively, reconciling the higher EU incorporation with the reduced formation of the 45S/47S rRNA precursors observed in FANCA-depleted cells. Transcription efficiency largely relies on the presence of G-quadruplexes (G4s), secondary DNA structures particularly enriched in rDNA coding sequences where their presence facilitates and optimizes RNAPolI progression (Cammas and Millevoi, 2017; Hall et al., 2017; Rhodes and Lipps, 2015; Richard and Manley, 2016; Santos-Pereira and Aguilera, 2015). Thus, we monitored by immunofluorescence the level of G4s in the nucleoli of FANCA-and FANCD2-deficient cells and observed that FANCA deficiency was associated with lower levels of G4s than in FANC pathway-proficient cells (Fig. 4G and F). In contrast, FANCD2 loss resulted in an increased level of nucleolar G4s (Fig. 4F).

It is generally assumed that transcription rate is negatively regulated by R-loops but fostered by G4 structures. Thus, in light of our data, it is tempting to speculate that in FANCD2-depleted cells the observed G4s increase compensates for R-loops increase resulting in unperturbed rDNA transcription. In contrast, FANCA depletion is associated to both R-loops increase and G4s decrease, a situation that possibly leads to the observed downregulation of the rDNA transcription. The reduced levels of nucleolar G4s in FANCA-depleted cells could be due to the more robust delocalization of NCL and NPM1, recognized as major determinants of the G4s stability inside the nucleolus (Cammas and Millevoi, 2017; Rhodes and Lipps, 2015).

### FANCA deficiency leads to altered ribosome profile and reduced protein synthesis

Several of our previous observations undoubtedly pinpoint that FANCA loss-of-function affects nucleolar homeostasis more deeply than FANCD2 or the other FANCcore complex partners. FANCA depletion deregulated nucleolar proteins localization not only in S/G2 but also in G1, affects rDNA transcription and increases the frequency of FANCC-or FANCG-mutated cells with nucleolar abnormalities. It is therefore tempting to speculate that, independently of the other partners of the FANCcore complex, FANCA could encroach on other cellular functions whose perturbations could be involved in the increased level of nucleolar stress observed in FANCA-depleted cells. In light to above, we decided to push our analysis further to understand why nucleolar homeostasis appears more pronounced in absence of FANCA. We failed to co-immunoprecipitate endogenous FANCA with the WT-NPM1, UBF, FBL or NCL and not any interesting candidate was identified in a yeast two-hybrid screen (data not shown), which is not in favour of a direct, interaction-mediated, role of FANCA in the maintenance of a nucleolar protein inside the nucleolus. Immunofluorescence analysis in cells transiently expressing a YFP-FANCA construct showed a nucleolar occurrence of FANCA and its enrichment in the granular compartment (GC), where NPM1 is also localized and the place where the pre-ribosomes are assembled (Fig. 5A). So, we reasoned that FANCA depletion could affect the pre-ribosome assembling, transport, processing, interaction with mRNA and/or the translation process, which alterations, in turn, may contribute to the loss of nucleolar homeostasis. To determine if FANCA-depleted cells are undergoing ribosome biogenesis abnormalities, we analysed their translation efficiency and their ribosome profile. Nascent proteins synthesis was determined by OP-Puro incorporation in FANCA^−/-^ (HSC72), FANCC^−/-^ (HSC536) lymphoblasts, their corrected counterpart as well as in FANC pathway-proficient cells (HSC93 and GM3657). Notably, only FANCA mutated lymphoblasts showed a consistent downregulation of proteins synthesis (Fig. 5B to D), also validated in FANCA-deleted HeLa cells (Fig. 5E). Given the previous data, we looked at the distribution of ribosomes on the mRNAs during their translation. The profiling revealed higher ratio monosomes vs polysomes in FANCA-deficient cells than in their corrected counterpart or in FANCA-proficient cells (Figure 5F and G).

**Figure 5:**
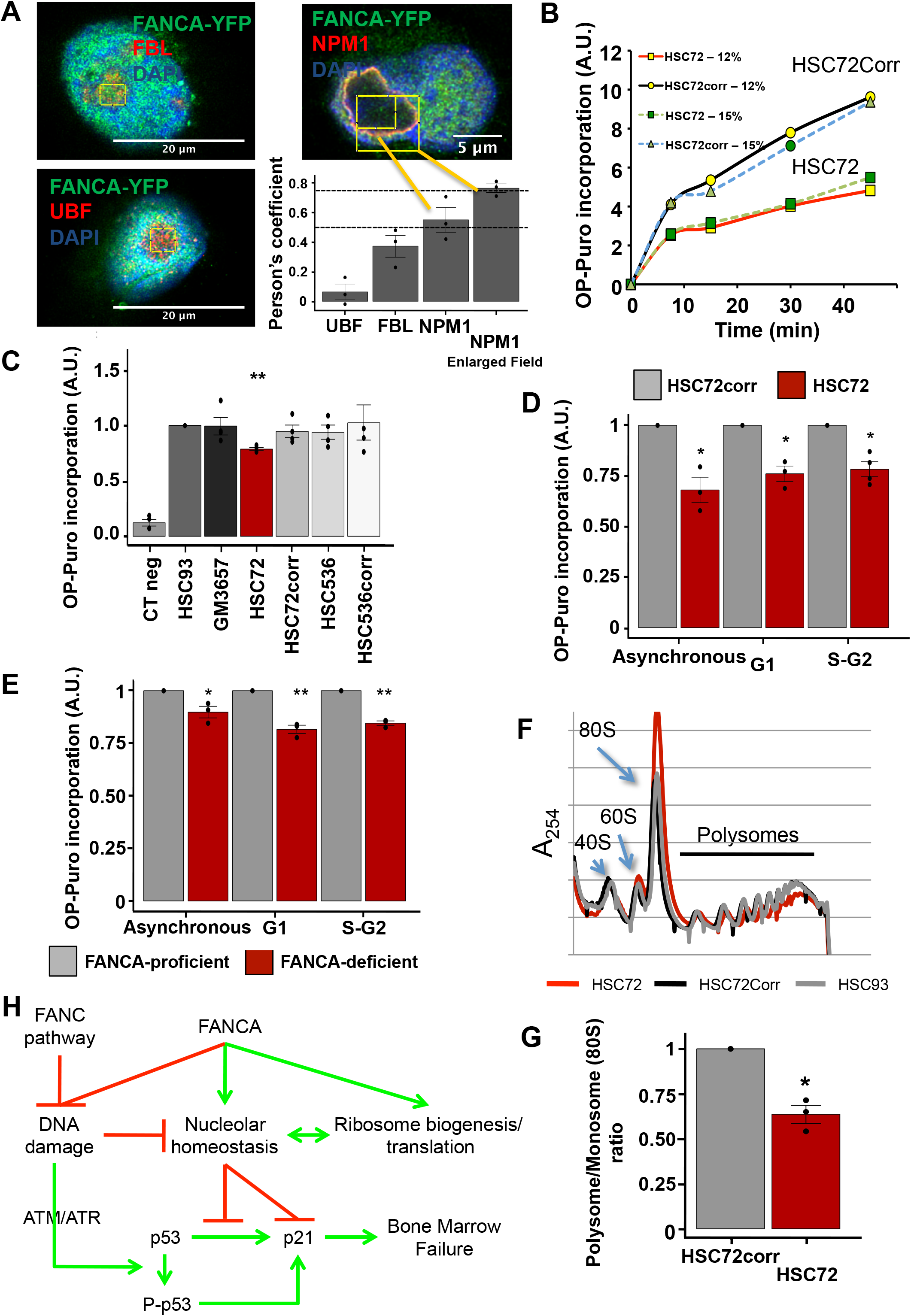
FANCA deficiency leads to altered ribosome profile and reduced protein synthesis. **A.** Cellular localization of FANCA as evaluated by confocal microscopy analysis of fixed cells transiently expressing a YFP-tagged FANCA construct. Cells were counterstained with FBL, UBF, or NPM1 to identify nucleoli and DAPI to stain DNA. JACoP analysis illustrates a significant colocalization of FANCA only with NPM1. **B.** Representative experiment showing a time-course incorporation of OP-Puro, i.e. nascent protein synthesis, assessed by FACS in exponentially growing lymphoblasts from a FANCA patient (HSC72) and their corrected counterpart (HSC72corr) cultured in 12% or 15% FCS. **C.** Diagram presenting the relative levels of OP-Puro incorporation in FANC-proficient (HSC93 and GM3657), FANCA^−/-^ (HSC72) and FANCC^−/-^ (HSC536) or FANC-corrected lymphoblasts. Bars represent the mean of 3 independent experiments. The value observed in HSC93 cells was settled at 1 in each individual experiment. Statistical significance was assessed with two-tailed Student *t*-test (**P<0.01). **D.** Histogram presenting the relative levels of OP-Puro incorporation in lymphoblasts from a FANCA patient (HSC72) and their corrected counterpart (HSC72corr) as a function of the cell cycle phase of the cell determined by staining with an anti-CyclinA2 antibody. The value of the HSC72corr cells was set to 1 in each individual experiment. Statistical significance was assessed with unpaired two-tailed Student’s *t*-test (*P<0.05). **E.** Diagram presenting the relative levels of OP-Puro incorporation in HeLa cells 72h after transfection, as function of the cell cycle phase. Bars represent the mean of 3 independent experiments. The value observed in FANCA-proficient cells was settled to 1 in each individual experiment. Statistical significance was assessed with two-tailed Student *t*-test (*P<0.05; **P<0.01). **F.** Representative experiment of a polysome profiling in FANCA-deficient HSC72 lymphoblasts and FANCA-proficient, HSC72corr and HSC93, cell lines. **G.** Quantification of the polysome/monosome ratio in HSC72 relative to HSC72corr cells. Bars represent the mean of 4 independent experiments +/-sem. The value of the HSC72corr cell was set to 1 in each individual experiment. Statistical significance was assessed with two-tailed Student’s *t*-test (*P<0.05). **H.** Model resuming our observations.

The easier explanation of our observations is that, in absence of FANCA, monosomes are charged on mRNAs but they start or progress slowly, resulting in altered polysome profiles and low levels of translation, which in turn could contribute to the observed loss of nucleolar homeostasis. Several reasons could be evoked as responsible for the observed abnormalities in translation, including imbalances in ribosomal proteins, defects in pre-ribosome assembling, transport and maturation, imbalances in non-ribosomal factors involved in translational initiation, progression and termination, alterations in signalling pathways involved in translation regulation. Further, in deep, studies are required to identify the mechanism involved in the translational imbalance associated to FANCA loss-of-function.

## Conclusions

Given the observed mislocalization of nucleolar protein in FANC pathway-deficient cells, we propose that, in line with its canonical functions in avoiding DNA breakage and DNA rearrangements, the pathway preserves the nucleolar homeostasis by allowing the faithful and optimal replication the rDNA sequences. Since their loss-of-functions leads to accumulation of DNA breaks and rearrangements during DNA replication, the depletion of FANC proteins participate to the observed transient nucleolar proteins mislocalization in S/G2 without causing major abnormalities in rDNA transcription. Our observations also revealed that the elevated expression of p21 in FA, considered as the key pathological event leading to the BMF in patients, depends on both DNA damage and nucleolar stress, in p53-dependent and -independent manner (Fig. 5H). In the context of the clinical phenotype of the FA syndrome, we observed that FANC pathway loss-of-function induces a redistribution of NPM1 outside the nucleolus, a known consequence associated to mutations in *NPM1*, which represent the most frequent mutagenic event in sporadic AML. Indeed, mutated NPM1 accumulates into the cytoplasm (Falini et al., 2005; Meani and Alcalay, 2009), where it can be immunoprecipitated with FANCA or FANCC (Du et al., 2010). Thus, the observed relocalization of NPM1 outside the nucleolus of FA cells could participate to the pre-leukemic status of the FA patients.

In addition, we unveiled a new, FANC pathway independent, role of FANCA demonstrating that its loss-of-function is associated to defects in rDNA transcription and mRNA translation that we failed to observe in other FANC pathway-deficient cells. These observations strengthen several previous observations demonstrating specific roles of the individual FANC proteins, and FANCA in particular, regardless of their participation to the FANC pathway (Adachi et al., 2002; Benitez et al., 2018; Nepal et al., 2018).

Finally, our observations drive FA closer to the group of iBMF syndromes called ribosomopathies that includes DC, DBA and SDS, in which BMF is associated with mutations in protein coding genes which loss-of-function directly leads to alteration in ribosome biogenesis and nucleolar homeostasis, supporting the centrality of the nucleolar metabolism in the maintenance of the bone marrow functionality.

## Supporting information

Supplementary informations and Figures

## Acknowledgements

A.G. was a fellow of the INSERM School Liliane Bettencourt and was supported by a “Course of Excellence in Oncology – Fondation Philanthropia” award. Work was funded by a grant from the La Ligue Contre Le Cancer to FR. The authors thank all the members of the UMR8200-CNRS research unit for helpful discussions; the Imaging and Cytometry Platform, UMS 23/3655, Gustave Roussy Cancer Campus, Univ. Paris Saclay, Villejuif, France; P. Pasero for the gift of the S9.6 hybridoma-purified antibody and for sharing protocols; KJ Patel for the gift of the FANCA-YFP plasmid; the Fanconi Anemia Research Fund (FARF) for the gift of antibodies; and A. Londono for the gift of the G4 antibody.

## Author contributions

A.G., G. R., and S.S.B.: Performed experiments. A.G. and F.R: Conceived and designed experiments. S.A. and J.J.D: Provided expertise on nucleolar metabolism and ribosome biogenesis. A.G., S.A., J.J.D. and F.R.: Analysed the data. A.G. and F.R.: Wrote the paper. F.R: Financed the study.

