## Supplementary informations and Figures for "Fanconi anemia proteins are required to maintain nucleolar homeostasis"

Material & Methods
Table 1 to 5, related to Material and Methods section
Supplementary Figure 1 & Legend
Supplementary Figure 2 & Legend

### Material & Methods

**Cell lines, culture and reagents:** Cell lines were listed in Table 1. HeLa, U2OS and HCT116 cells and human and mouse primary fibroblasts were grown in Dulbecco's modified Eagle's medium (DMEM) (Life Technologies) supplemented with 10% FCS, 0.5 mg/ml penicillin, 100 µg/mL streptomycin and 1 mM pyruvate, at 37°C and 5% $CO_2$. Human lymphoblasts were grown in RPMI medium (Life Technology) supplemented with 12% FCS, 0.5 mg/ml penicillin, 100 µg/mL streptomycin and at 37°C and 5% $CO_2$. Cell cultures were regularly tested for the absence of mycoplasma.

ATM and/or ATR inhibitors, caffeine 1 mM (Sigma), KU-55933 10 µM (Abcam) and VE-822 100 nM (Sellchem), were added to culture medium 6h after transfection. The addition was repeated every 24h until the end of the experiment. Actinomycin D (Sigma) was used at 10 ng/mL for 16h.

**siRNA and plasmid transfection:** siRNA sequences are listed in Table 2. siRNAs were transfected at 20 nM using calcium phosphate diluted at 6.25 mM in HBSP buffer (750 µM $Na_2HPO_4$, 5 mM KCl, 140 mM NaCl, 25 mM HEPES pH 7). FANCA-YFP plasmid (a gift from KJ Patel) was transfected using jetPRIME (Polyplus) in HeLa cells or Fugene HD (Promega) in U2OS cells.

**Western blotting:** Antibodies are listed in Table 3. Cells were lysed in a 50 mM Tris-HCl pH 7.5, 125 mM NaCl, 1 mM MgCl, and 0.1% SDS buffer supplemented with benzonase (Merck, 1:1000), phosphatase inhibitor PhosSTOP (Roche) and cOmplete ULTRA EDTA-Free Protease inhibitors (Roche). Proteins were resuspended in Laemmli's blue (16 mM Tris-HCl pH 6.8, 10% glycerol, 2.5% SDS) supplemented with β-mercaptoethanol (Sigma, 1:10). Migration was performed at 150V in a 2.5 mM Tris, 19.2 mM glycine, and 0.01% SDS buffer. Semi-dry transfer was performed with a TransBlot cell apparatus (Bio-Rad) in transfer buffer composed of 2.5 mM Tris-HCl pH 8.3, 19.2 mM glycine, 0.01% SDS, and 20% isopropanol for 1h at 20V. Liquid transfer was performed in an XCell II module (Invitrogen) in a 2.5 mM Tris-HCl pH 8.3, 19.2 mM glycine, 0.04% SDS, and 20% methanol buffer at 15 V O.N. at 4°C. Nitrocellulose membranes (Protran 0.2 mm, Amersham) were blocked for at least 30 min in 0.1% PBS-Tween, 5% milk and incubated with primary antibodies in TBS/0.1% Tween/5% BSA or PBS/0.1% Tween/5% milk. Visualisation was performed using ECL (Life), Western Bright ECL (Advensta) or Immobilon (Millipore) developer. Images were acquired using a CCD camera (General Electrics).

**Immunofluorescence and image quantification:** Antibodies are listed in Table 3. Samples were fixed for 48h to 72h after siRNA transfection. For FBL/NCL/NPM1, cells were washed with PBS, fixed in 3.9% formaldehyde (Sigma) for 5min at R.T. and permeabilised in PBS with 0.5% Triton (Sigma) for5min at R.T. For UBF, cells were fixed in 3.9% formaldehyde (Sigma) for 5 min at R.T. and permeabilised in PBS/0.5% Triton (Sigma) for 5min at R.T. For R-loops and G-quadruplex, cells were washed with PBS, fixed in methanol at -20°C for 5 min, then blocked in PBS, 0.05% Tween, and 3% BSA for 1h at R.T. and stained with primary followed by secondary antibodies in the same buffer. Epifluorescent images were acquired with a Zeiss microscope at X63 magnification 1.4 ON. Quantification was performed on at least 100 cells. Nucleolar and nucleoplasmic intensity quantification of R-loops and G-quadruplexes was performed on epifluorescence images using CellProfiler software and using fibrillarin staining as a nucleolar marker. For colocalisation assessment, image acquisition was performed using a confocal Leica TCS Sp8 microscope at X63 magnification 1.4 ON (with magnification to a resolution of approx. 40 nm/pixel). Deconvolution was performed using Huygens software. Colocalisation assessment was performed using the JACoP plug-in on ImageJ software. A negative control was performed acquiring each signal separately.

**Electronic microscopy analysis**: Ultrastructural study: 72h after siRNA transfection, cell monolayers were fixed in 2% glutaraldehyde in 0.1 M Sörensen phosphate buffer for 1h at 4°C. The cells were scraped off during fixation and centrifuged. The fixed pellets were rinsed for 1 h in ice-cold phosphate buffer, post-fixed with 2% aqueous osmium tetroxide and dehydrated in increasing concentrations of ethanol prior to Epon embedding. Polymerisation was carried out for 48h at 60°C. Ultrathin sections were stained with standard uranyl acetate and lead citrate prior to observation with an FEI Technai Spirit transmission electron microscope at 80 Kv. Digital images were taken with an SIS MegaviewIII CCD camera.

**EdU, EU, and OP-Puro labelling:** EdU was purchased from Thermo Fisher Scientific, as part of the Click-iT® EdU Alexa Fluor® 488 Imaging Kit (#C10337, Thermo Fisher Scientific). EU (#BCN-003-5, BaseClick) and O-Propargyl-puromycin (#NU-931-05, Jena Bioscience) were from the same kit using 5-FAM (Lumiprobe) as a fluorophore. 72h after transfection, labelling was performed by adding EdU (10mM) 10min; EU (1mM) 15min; OP-Puro (50mM) 15 min. Labelling was analyzed by flow cytometry (C6, BD Accuri apparatus).

**Metabolic labelling:** Seventy-two hours after siRNA transfection, non-confluent HeLa cells were incubated for 1h at 37°C and 5% $CO_2$ in phosphate-free DMEM medium containing 10% dialysed FCS. They were then pulsed for 20min with 15 mCi of ($^{32}$P)orthophosphate (Perkin Elmer). After 2 washes in culture medium, they were incubated for a chase period of 1 to 6 h. RNAs were extracted using TRIzol reagent, and 1 mg of RNA was loaded on a denaturing gel with BET. The BET signal was acquired after migration. The gel was dried, and the radioactive signal was acquired using Typhoon FLA9500 scanner (GE Healthcare).

**Northern blot:** Seventy-two hours after siRNA transfection, 1 mg of TRIzol-extracted RNA was loaded on a denaturing gel with BET. RNA was passively transferred on a nitrocellulose membrane (Amersham) in SSC 10X buffer (1.5 M NaCl, 0.15 M sodium citrate, pH 7). The BET signal was acquired after transfer. Membranes were blocked in 3X SSC, 5X Denhardt's, 0.5% SDS, and 0.5 mg/ml ARNt for 1 h at 45°C. Northern blot probes (described in Supplemental Table 3) were labelled with 25 mCi ($^{32}$P)ATP (Perkin Elmer) using T4 polynucleotidekinase (NEB). Incubation was performed O.N. at 45°C. Radioactive signal was acquired using Typhoon FLA9500 scanner (GE Healthcare).

**ChIP:** HeLa cells (10-15x10$^6$) were grown in Petri dishes (confluency approx. 70-80%). Forty-eight hours after transfection, medium was replaced by fresh medium, and fixation was performed as recommended by the ChIP-IT High sensitivity kit (Active Motif) (15 min in 1.1% PFA fixation). Cells were then processed using the ChIP-IT High sensitivity kit, and immunoprecipitation was performed using 4 µg of antibodies listed in Table 3.

**ChIP-qPCR:** Probes are listed in Table 5. For ribosomal DNA ChIP-Q-PCR, the following programme was used: 50°C, 2min; 95°C, 10min; 45x (95°C, 15s; 58°C, 1min/30s); dissociation stage (95°C, 15s; 60°C, 1min; 95°C, 15s; 60°C, 15s)..

**PCR-based analysis of rDNA rearrangement:** 72h after transfection, cells were lysed and DNA was extracted using the Maxwell RSC Blood DNA kit (Promega). PCR was performed on 100ng of DNA using 0.02U/mL of Phusion polymerase, 0,2mM of dNTP and 0.5mM of the followig probes: F: 5'AGTCGGGTTGCTTGGGAATGC3', R: 5'GGACAAACCCTTGTGTCGAGG3' (predicted position in gene: nt 8204-12970). The following amplification programme was used: 98°C, 30s; 35x (98°C, 10s; 62°C, 30s; 72°C, 4min); 72°C, 10min.

**Proximity Ligase Assay (PLA):** Seventy-two hours after siRNA transfection, cells were fixed, and PLA was performed using DUOLINK (Sigma) reactive molecules according to manufacturer's recommendations. Antibody dilutions were as follows: in the HCT-WT cell line (p21: 1/2000, NPM1: 1/2500), in the HCT-p53$^{-/-}$ cell line (p21: 1/200, NPM1: 1/2000).

**Polysomal fractionation:** Cycloheximide (100µg/ml) was added for5 min to cell cultures and was maintained in PBS washes. Cells were fractionated in 5mM Tris pH7.5, 2.5mM $MgCl_2$, and 1.5mM KCl buffer supplemented with cOmplete ULTRA EDTA-Free Protease inhibitors (Roche), 100µg/ml cycloheximide, RNase Inhibitor (Promega), 2mM DTT, 0.5% Triton and 0.5% sodium deoxycholate, and nuclei were discarded after a 7min, 16,000g centrifugation. The cytoplasmic OD at 260nm was assessed, and the lysate was diluted up to 10 to 20 OD. Then, 500µL was loaded on a 5-50% sucrose gradient (20mM HEPES pH7.6, 100mM KCl, 5mM $MgCl_2$, 10µM cycloheximide, 1/10 protease inhibitors, 10U/mL RNase inhibitor), and centrifuged for 2h at 36,000rpm at 4°C in a Beckman SW41Ti rotor. The A254 of the fractions was determined using a UA-6 UV/VIS Detector.

#### Table 1: list of the cell lines used

| Cell line | Origin | |
|---|---|---|
| HeLa | in house & ATCC | |
| U2OS | in house and ATCC | |
| HCT116 | Gift of M. Debatisse | |
| HCT116-p53$^{-/-}$ | Gift of M. Debatisse | |
| HSC93 | in house | Gift of M. Buchwald lab, received in 1990 |
| HSC72 | in house | Gift of M. Buchwald lab, received in 1990 |
| HSC536 | in house | Gift of M. Buchwald lab, received in 1990 |
| HSC72CORR | in house | Gift of M. Buchwald lab, received in 1990 |
| HSC536CORR | in house | Gift of M. Buchwald lab, received in 1990 |
| GM3657 | Coriell repository | |
| GM3348 | Coriell repository | |
| GM3652 | Coriell repository | |
| GM5757 | Coriell repository | |
| GM16754 | Coriell repository | |
| GM00449 | Coriell repository | |
| GM02641 | Coriell repository | |
| GM13136 | Coriell repository | |
| GM16635 | Coriell repository | |
| MRC5 | in house | |
| MRC5-SV | in house | |
| AS911 | Gift of A. Sarasin lab | |
| FA Human Primary fibroblasts | Gift of J Soulier lab | Donor's name blided. |
| FA Mouse Primary fibroblasts | Derived in house | |

**Table 2: list of siRNA used**

| GENE | SEQUENCE |
|---|---|
| LACZ | CGUCGACGGAAUACUUCGA |
| FANCA | |
| FANCA1 | GUACAGCAGCAAUUUCUUA |
| FANCA2 | GAUCGUGGCUCUUCAGGAA |
| FANCA3 | GGACAUCACUGCCCACUUC |
| FANCD2 | |
| FANCD2_1 | GGAGAUUGAUGGUCUACUA |
| FANCD2_2 | AACAGCCAUGGAUACACUUGATT |
| FANCD2_3 | CAGAGUUUGCUUCACUCUCUA |
| FANCC | |
| FANCC_1 | GAGAGAAUCAUCUUAAUGG |
| FANCC_2 | GGAAUCGUCUUGGCAUUGA |
| FANCC_3 | GGUAUGCACCUAUAGAUUA |
| ATM | ATM1-492 (Invitrogen) |
| ATR | HSS100876, HSS10877, HSS878 (Invitrogen) |

### Table 3: list of antibodies used:

| Antibody target | Supplier | Code N° | Specie | WB dilution | IF dilution | CHIP | FACS |
|---|---|---|---|---|---|---|---|
| Alexa-Fluor Secondary Antibodies | Life Technologies | | D | | 1:1000 | | 1:150 |
| BrdU (Bu20a) | Dako | M0744 | M | | | | 1:60 |
| Cyclin A2 | Abcam | ab16726 | M | | 1:200 | | 1:100 |
| FANCA | Bethyl | A301-980A | R | 1:1000 | | | |
| FANCA | FARF | FANCA-1 | R | 1:1000 | | | |
| FANCC | FARF | FANCC-2 | R | 1:1000 | | | |
| FANCD2 | Novus | NB100-182 | R | | | 4µg | |
| FANCD2 (FI17) | Santa Cruz | sc-20022 | M | 1:1000 | | | |
| Fibrillarin | Abcam | Ab5821 | R | 1:1000 | 1:2000 | | |
| G-Quadruplexes (clone1H6) | Merck | MABE | M | | 1:200 | | |
| H2AX | Abcam | ab11175 | R | 1:1000 | | | |
| γH2AX (JBW301) | Millipore | 05-636 | M | 1:1000 | 1:2000 | | |
| H3K9me3 (6F12-H4) | Millipore | 05-12432 | | | | 4µg | |
| IgG1 | Dako | X0931 | M | | | 4µg | |
| Lamin | Santa Cruz | sc-7292 | M | 1:1000 | | | |
| NPM1 (FC61991) | Thermo Fisher | 32-5200 | M | 1:1000 | 1:500 | | |
| Nucleolin (4E2) | Abcam | ab13541 | M | | 1:4000 | | |
| p21 (12D1) | Cell Signaling | 2947 | R | 1:1000 | | | |
| p53(DO-7) | Santa Cruz | sc-47698 | M | 1:1000 | | | |
| p53Ser15 | Cell Signaling | 9284S | R | 1:1000 | | | |
| RNAPolI | Abcam | ab101977 | | | | | |
| S9.6 | P. Pasero's team | | M | | 1:200 | | |
| UBF (F9) | Santa Cruz | sc-13125 | M | 1:1000 | 1:250 | 4µg | |
| Vinculin (spm227) | Abcam | ab-18058 | M | 1:4000 | | | |

FARF: Fanconi Anemia Reseaech Found; R: Rat; M: Mouse; D: Donkey

### Table 4: Northern blot probe used

| Name | Sequence | Position in gene<br>Genbank accession #U13369 |
|---|---|---|
| **5'ETS1b** | 5'-AGACGAGAACGCCTGACACGCACGGCAC-3' | 297 to 324 |

### Table 5: list of the ChIP-qPCR primers used

| Name | Forward | Reverse | Efficiency $(10^{(-1/slope)}-1)$ | Elongation T° | Position | Ref. |
|---|---|---|---|---|---|---|
| H1 | GGCGGTTTGAGTGAGACGAGA | ACGTGCGCTCA2CCGAGAGCAG | 0,95 | 58° | 952-1030 | Nat. Cell Biol. 7, 311–318 (2005) |
| H8 | AGTCGGGTTGCTTGGGAATGC | CCCTTACGGTACTTGTTGACT | 0,90 | 58° | 8204-8300 | Nat. Cell Biol. 7, 311–318 (2005) |
| H27 | CCTTCCACGAGAGTGAGAAGCG | CTCGACCTCCCGAAATCGTACA | 1,11 | 58° | 27366-27477 | Nat. Cell Biol. 7, 311–318 (2005) |
| H42.9 | CCCGGGGGAGGTATATCTTT | CCAACCTCTCCGACGACA | 1,05 | 58° | | Nat. Cell Biol. 7, 311–318 (2005) |

**Extended Data Figure Legends:**

**Figure S1:**

**A.** Western blot showing the consequence of FANCA or FANCD2 siRNA-mediated depletion in HeLa or U2OS cells 72h after transfection on FANCA, FANCD2, NPM1, NCL and FBL expression. Vinculin was utilized as loading control.

**B.** Confocal microscopy images showing wide fields of HeLa cells after transfection with untargeted (siLacZ) or FANCA-targeted siRNAs stained with antibodies against the nucleolar proteins UBF (Red) and FBL (Green), and counterstained with DAPI to visualize DNA

**C.** Immunofluorescence microscopy images showing wide fields of HeLa cells 72h after transfection with untargeted (siLacZ) or one of the three siRNA, siFANCA1 to siFANCA3, that pooled were used in Fig.1a. Cell were stained with antibodies against the nucleolar proteins NCL (Red) and FBL (Green) and counterstained with DAPI, to visualize DNA.

**D.** Percentage of cells with altered nucleoli following transfection with each individual FANCA targeted siRNA sequence, siFANCA1 to siFANCA3, otherwise pooled into the siFANCA. In the box, WB illustrating the shutdown of FANCA expression induced by each single siRNA is shown. Vinculin was utilized as loading control. Bars represent the means of 3 independent experiments +/- sem.

**E.** Percentage of cells showing 1, 2 or more nucleoli as a function of FANCA expression in HeLa (Left panel) or U2OS (Right panel) cells. Bars represent the means of 3 independent experiments +/- sem.

**F.** Immunofluorescence microscopy images showing single nuclei of primary fibroblasts from 3 to 5 months old WT or Fanca$^{-/-}$ mice stained as described above.

**G.** Electronic micrographs showing nucleolar morphology as observed in primary from WT (i) or Fanca$^{-/-}$ mice fibroblasts (ii and iii).

**H.** Percentage of primary fibroblasts presenting canonical nucleolar morphology (irregular shape) or a round or cap-like shape. Analyses of fibroblasts isolated from two WT and two Fanca$^{-/-}$ mice are reported.

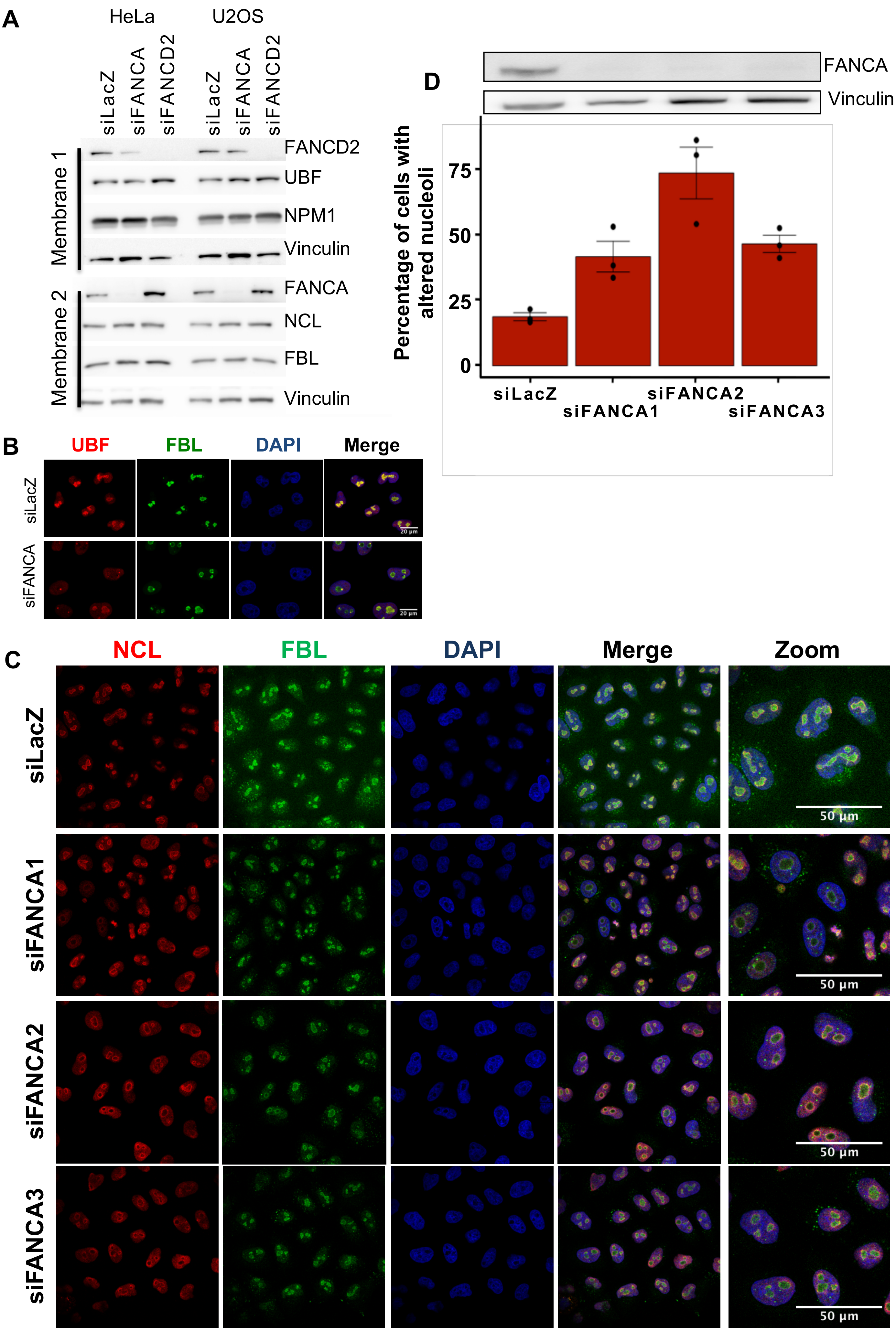
A
HeLa
U2OS
siLacZ
siFANCA
siFANCD2
Membrane 1
Membrane 2
FANCD2
UBF
NPM1
Vinculin
FANCA
NCL
FBL
B
UBF
FBL
DAPI
Merge
20 µm
C
NCL
Zoom
siFANCA1
siFANCA2
siFANCA3
50 µm
D
Percentage of cells with altered nucleoli
0
25
50
75

E

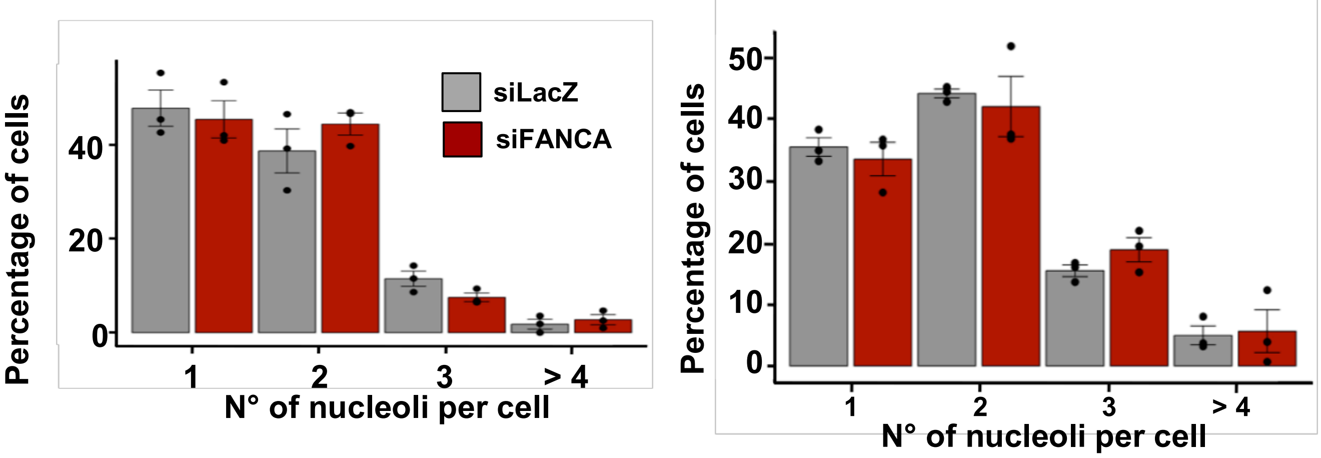

Percentage of cells
40
20
0
1
2
3
> 4
N° of nucleoli per cell
siLacZ
siFANCA
Percentage of cells
50
40
30
20
10
0
1
2
3
> 4
N° of nucleoli per cell

F

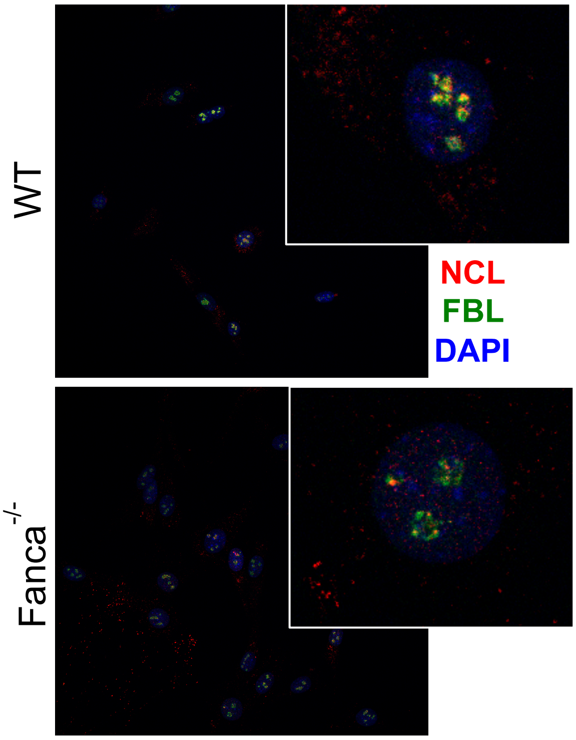

WT
Fanca-/-
NCL
FBL
DAPI

G

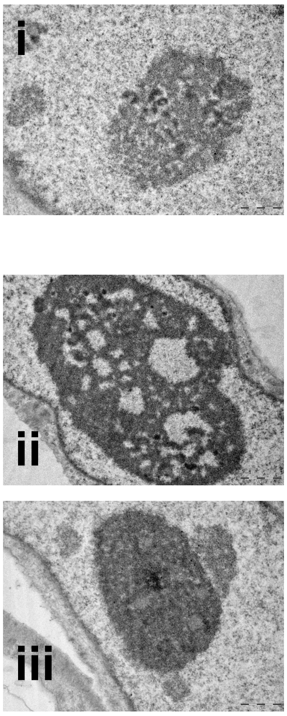

i
ii
iii

H

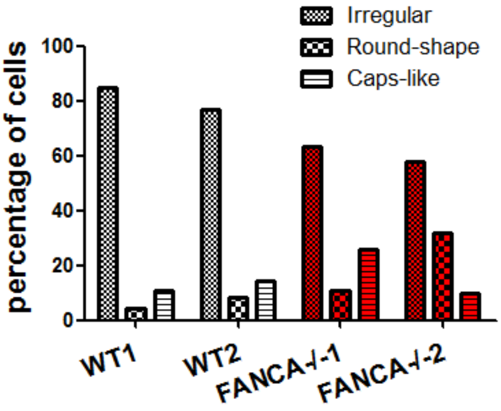

Irregular
Round-shape
Caps-like
percentage of cells
100
80
60
40
20
0
WT1
WT2
FANCA-/-1
FANCA-/-2

**Figure S2.**

**A and B.** Dots in the diagram represent the intensity of R-loop-staining measured on epifluorescence images using CellProfiler software with FBL staining as nucleolar marker and DAPI staining as marker for the nucleus surface for each individual HeLa cell 72h after transfection with the indicated siRNAs. Cells were incubated with the RNaseH that, targeting specifically the DNA-RNA hybrids, validates antibody specificity. A representative experiment is reported. At least 100 cells were scored for each condition. Statistical significance was assessed with unpaired two-tailed Student *t*-test (***P<0.005).

**C.** Western blots showing the expression of the indicated proteins in HCT116 or HCT116-p53$^{-/-}$ cells transfected with untargeted (siLacZ) or/and targeted (FANCA or/and NPM1) siRNAs.

**D.** 72h following transfection with the indicated siRNAs**,** HeLa cells were labeled 20min with ($^{32}$P)orthophosphate and chased with cold orthophosphate for 1, 2, 3 and 6h. Size of the processing intermediates and mature rRNA species are indicated. Ethidium bromide (EtBr)-stained gel is shown in the bottom as a loading control. A representative experiment is showed.

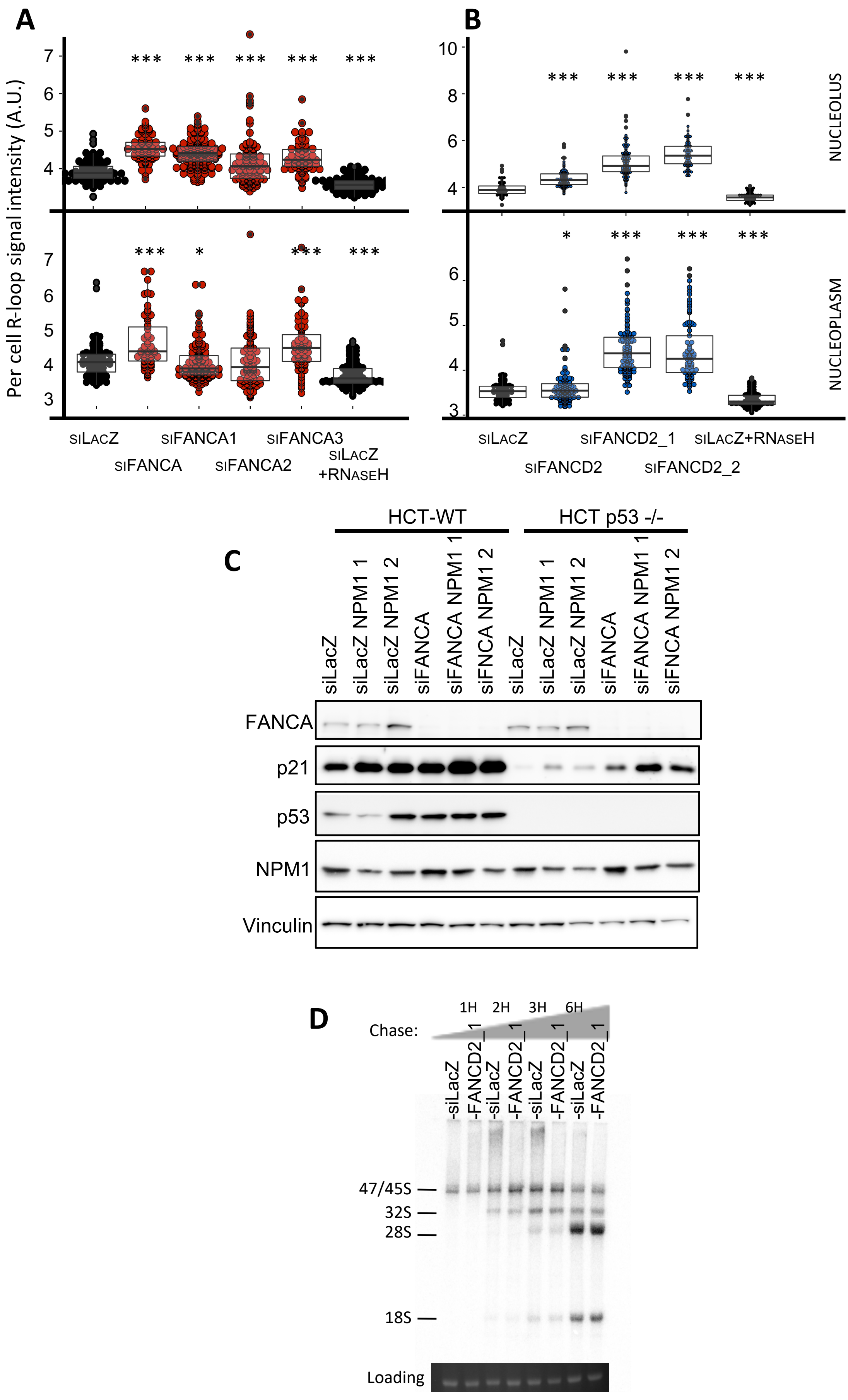

A
B
C
D
Per cell R-loop signal intensity (A.U.)
NUCLEOLUS
NUCLEOPLASM
siLacZ
siFANCA
siFANCA1
siFANCA2
siFANCA3
siLacZ +RNaseH
siFANCD2
siFANCD2_1
siFANCD2_2
siLacZ+RNaseH
HCT-WT
HCT p53 -/-
siLacZ
siLacZ NPM1 1
siLacZ NPM1 2
siFANCA
siFANCA NPM1 1
siFNCA NPM1 2
FANCA
p21
p53
NPM1
Vinculin
Chase:
1H
2H
3H
6H
-siLacZ
-FANCD2_1
47/45S
32S
28S
18S
Loading
